# Omada: Robust clustering of transcriptomes through multiple testing

**DOI:** 10.1101/2022.12.19.519427

**Authors:** Sokratis Kariotis, Tan Pei Fang, Haiping Lu, Chris Rhodes, Martin Wilkins, Allan Lawrie, Dennis Wang

## Abstract

Cohort studies increasingly collect biosamples for molecular profiling and are observing molecular heterogeneity. High throughput RNA sequencing is providing large datasets capable of reflecting disease mechanisms. Clustering approaches have produced a number of tools to help dissect complex heterogeneous datasets, however, selecting the appropriate method and parameters to perform exploratory clustering analysis of transcriptomic data requires deep understanding of machine learning and extensive computational experimentation. Tools that assist with such decisions without prior field knowledge are nonexistent. To address this we have developed Omada, a suite of tools aiming to automate these processes and make robust unsupervised clustering of transcriptomic data more accessible through automated machine learning based functions. The efficiency of each tool was tested with five datasets characterised by different expression signal strengths to capture a wide spectrum of RNA expression datasets. Our toolkit’s decisions reflected the real number of stable partitions in datasets where the subgroups are discernible. Within datasets with less clear biological distinctions, our tools either formed stable subgroups with different expression profiles and robust clinical associations or revealed signs of problematic data such as biased measurements.

## Introduction

The rapid development of next-generation sequencing boosted the quantitative analysis of gene expression in a variety of human tissues and organs^1,2^ generating valuable resources^3^ for downstream investigative analysis. In recent years, such analyses aim to elucidate disease mechanisms^4^ and construct genomic profiles^5^ to explain diagnosis^6^, prognosis and treatment patterns. However, transcriptomic profiles can be heterogeneous due to several causes pertaining to technical biases that produce batch effects^7^, cellular diversity^8^, disease heterogeneity^9^ as well as differences between individuals and populations^10,11^. In turn, this heterogeneity hinders traditional research efforts^12^ aiming to define structures especially under complex diseases which led to the utilisation of the field of machine learning towards this data demanding goal. More specifically, unsupervised machine learning i.e. clustering, in the form of transcriptomic profiling based on sequencing data^13–15^, has been explored in terms of symptomatic heterogeneity in complex diseases revealing differences in molecular states^16–18^ and phenotypes described by the gene expression of diseased tissue. However, deep medical and molecular knowledge is required to identify solvable problems and interpret results within the context of various diseases and conditions. Simultaneously, specialised knowledge and experience is needed to create functional, efficient and insightful models which generate reproducible solutions. Despite the inherent power of these models, most times a default model is not sufficiently tuned to the specific dataset thus unable to extract essential information. Many models have been compared, tested and found to work on different data and research questions^19,20^ highlighting that no single model is always optimal without tuning (or optimising) on the specific dataset at hand, especially with state-of-the-art methodologies^21,22^.

Machine learning (ML) is currently being used in many forms and combinations^23^, for different types of projects within diverse fields of biomedical research^24–26^. Supervised and unsupervised methods are being developed to address specific questions and/or data problems as the pace of new data generation increases rapidly. Big data has made the importance of tailored methodologies essential for specialised datasets^27,28^, as speed and accuracy pose an even greater obstacle, especially when handling sizable medical data. The impact of machine learning in biomedical sciences has risen considerably with the multitude of methodologies leading to previously unfeasible computations^29,30^. Unsupervised learning proved to be an invaluable tool towards exploring heterogeneity in complex diseases since its functioning without any prior knowledge or assumptions of sample labels. Due to this diversity, there is a need for methods that support non-expert users to utilise the characteristics of various methodologies in their unsupervised work. One of the most important aspects of sample partitioning is the stability of the generated groups as unstable clusters, usually imply the lack of signal which should be present and drive the clusters. Signals can take many forms, for example the level of gene activity in RNA sequencing datasets. Cluster instability can be caused inherently by the data points or by the type and application quality of a clustering technique.

With the above obstacles in mind, we introduce Omada, a toolkit with multiple functions based on cluster stability and machine learning formulas to provide assistance to both experienced and inexperienced users during the steps from dataset assessment to the formation of the subgroups. Each function’s results are based on machine learning theory and multiple metrics to ensure that a wealth of methods will be considered in the decision and clustering process.

## Methods

This toolkit consists of a pipeline that takes in a gene expression matrix to identify transcriptomic subgroups of samples (Figure 1 and Supplementary Figure 1). Starting from a matrix of gene expression values (e.g. transcripts per million from RNAsequencing), the most suitable clustering method is chosen, followed by selecting the transcript features for clustering and determining the number of clusters and memberships.

**Figure 1:**
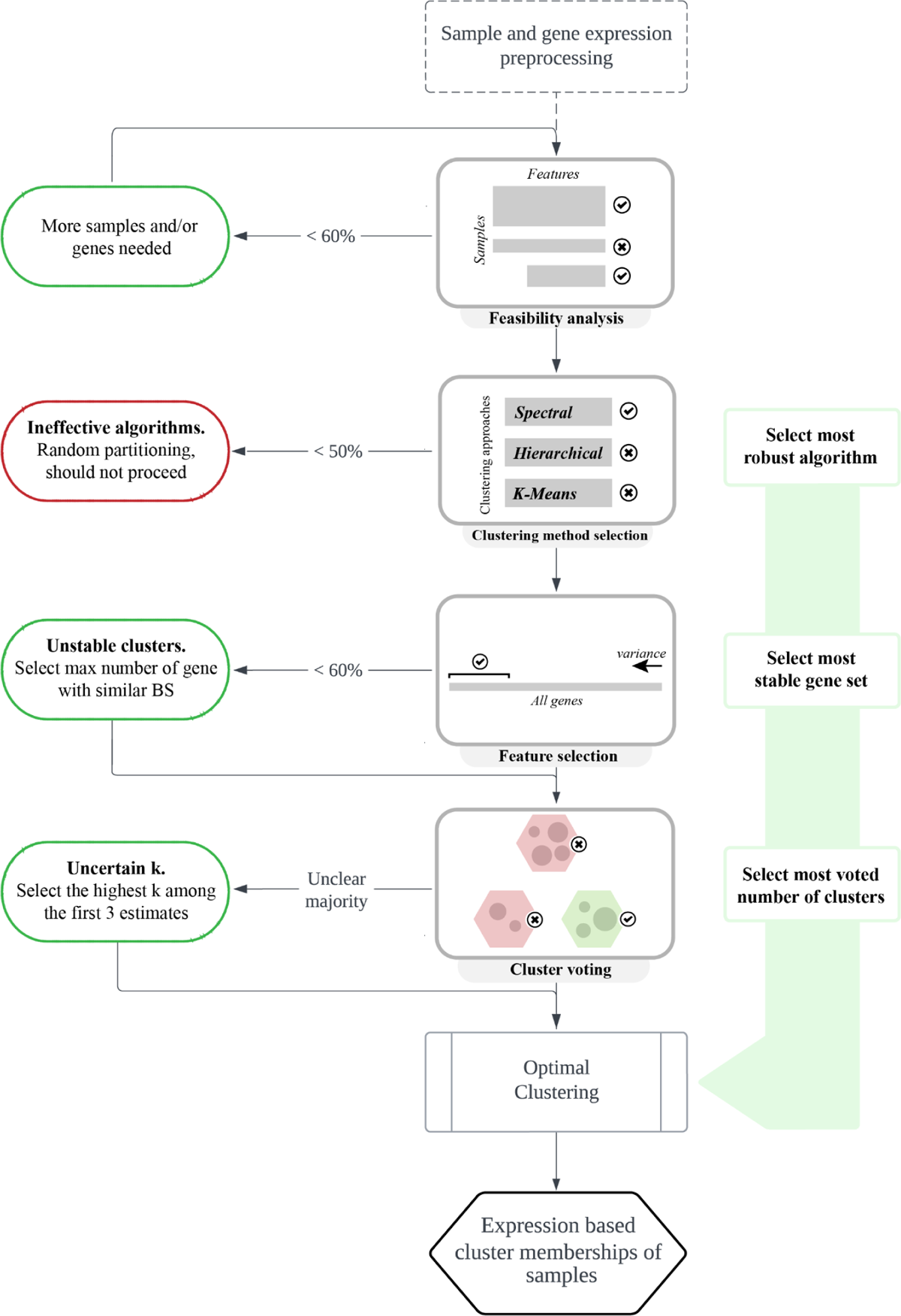
An overview of steps for discovering gene expression subgroups using Omada’s clustering tools. First, processing, quality control and feasibility analysis are ensuring the input data are suitable for clustering. Then, choose the most robust clustering methodology that provides the most consistent partitions. Next, determine the genes that are useful for discriminating samples and provide the most stable clusters. Finally, determine the number of subgroups that potentially exist in the cohort by selecting the number of clusters (k) supported by the majority of internal machine learning indexes. The end result, after the final optimised clustering, consists of the assignment of a cluster to each sample driven solely by its expression profile.

### Sample and gene expression preprocessing

A preprocessing step is recommended by the user before the application of these tools on any dataset to heighten the chances of any underlying important signal to be discovered. Data biases and format can often drive clustering attempts to focus on discriminating data points based solely, or mostly, on known information producing no new insights irrespectively of the method used^21,31^. To address this, it is recommended to attempt to remove/normalise any data points that might be introducing strong biases to allow the novel signal to be detected. Furthermore, numerical data may need to be normalised in order to account for potential misdirecting quantities (i.e. outliers) or specifically transformed to satisfy an algorithm’s input criteria. Data points or samples have to be filtered based on field knowledge to allow the data to answer specific scientific questions. Expression data should go through proper quality control depending on the manner of collection to identify outliers and remove unreliable datapoints. For microarrays it’s important to assess sample, hybridization and overall signal qualities along with signal comparability and potential biases. Array correlations through PCA and correlation plots should also be considered^32^. RNA sequencing experiments also produce data that need to be controlled for potential trimming of adapter sequences, low quality reads, uncalled based and contaminants by using a plethora of available tools^33,34^. Additionally, qPCR generated data should be checked for abnormal amplification, positive and negative control samples, control on PCR replicate variation and determine reference gene expression stability and deviating sample normalisation factors^35^. As for the number of genes, it is advised for larger genesets (>1000 genes) to filter down to the most variable ones before the application of any function as genes that do not vary across samples do not contribute towards identifying heterogeneity. Moreover, large genesets require increased computational power and extended runtime without adding any real value due to the large number of non useful genes. Lastly, it is important to note that technical artefacts, such as sampling location or machine specifications, may drive clustering causing the formation of very distinct clusters which can solely be attributed to relevant biases. It is very important for those cases to be identified and extracted insights should be disregarded as they do not reflect real signals or data trends.

### Determining clustering potential

At the start of each study, we assess the suitability of the input dataset for clustering to ensure general dataset attributes do not influence the process (Figure 1). The number of samples and features -i.e. genes-, as well as the balance of the two dimensions, directly affects the capabilities of clustering methods to handle the dataset. An inadequate number of samples does not provide enough training power^36^, while an overabundance of samples might clutter the provided information and confuse most methodologies^37^. Similarly, too few features can lead to weak clustering criteria and too many features might lead a methodology away from the features that can really differentiate between clusters of samples. Therefore, to estimate the feasibility of a clustering procedure on a specifically sized dataset we rely on measurable metrics of cluster quality, such as stability. Clusters of high stability denote both a partitionable dataset as well as a dataset-suitable methodology^38^. The feasibility score of any dataset is a function of both dimensions as well as the number of classes requested. As such, if too many or a single class is requested of a relatively small dataset the calculation will reflect low feasibility due to insufficient samples and/or features to form the desired classes.

### Simulating datasets

To assess the quality of the dataset to be used, our toolkit includes two functions for simulating datasets of different dimensionalities for stability assessment. We use those to understand the relation between the number of samples, genes and cluster sizes. The first function simulates datasets allowing for selecting the number of samples (*n*), genes (*m*) and clusters (*c*). Each cluster contains 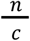 samples drawn from a normal distribution with a different mean and standard deviation. Each mean is drawn from a sequence of *c* evenly spaced integers that belong to the range [5, *c* * 10]. Each standard deviation is similarly drawn from a range of [1, *c* * 2]. To estimate the difference between distributions, we calculate the two sided Kolmogorov’s D statistic between each pair of distributions representing the generated classes and plot the empirical cumulative distribution function (EDCF).

Subsequently, we calculate the stability of each *k* (number of clusters for a particular clustering run) using the clusterboot function in R package fpc v2.2-3. The number of clusters *k* to be considered belong to *k* ∈ *[number of classes – 2, number of classes* + 2], with a minimum of *k* = 2. The maximum and average stabilities over all *k* are reported, providing a stability-based quality score that provides an insight in deciding whether a prospect dataset is suitable for a clustering study.

To assess the clustering feasibility of an existing dataset this tool kit also provides a similar function which generates a simulated dataset based on an input dataset and the user’s estimation of the number of classes. The number of samples and genes equal those of the input dataset and its mean (*m_input_*) and standard deviation affect those of each generated class within the dataset. Specifically, if *n* ∈ (1,2,3,…) is the number of classes, each class mean (*m_class_*) equals *m_input_* * 10 * *n* and each class standard deviation (*sd_class_*) equals *sd_input_* * 2 * *n*.

### Intra-method Clustering Agreement

Unsupervised learning offers a multitude of methods to be applied on specific types of data due to their nature (e.g. numeric, binary) or underlying signal to be detected. Most studies employ widely-used methods (e.g. hierarchical clustering) without exercising any kind of selection method that would point towards the most effective methodology. Selecting an appropriate approach requires extensive machine learning and data analysis knowledge coupled with tuning and testing of multiple different algorithms. To enable non-machine learning expert users to utilise the vast capabilities of this field and avoid default limited efficiency methodologies we present a clustering selection tool that offers an intelligent selection method with unbiased results through parameter randomization. The nature of this selection method allows any number of well established unsupervised methods to be considered.

To address the lack of class labels and thus a performance measure in unsupervised models, we compare how consistently different approaches partition our data when one or more parameters change. As high consistency we define the high agreement score calculated between different variations of a clustering algorithm. When two different clustering runs agree on the partitioning of the samples they also show robustness since they do not randomly assign samples to subgroups but rather are driven by the underlying structure of the data.

We implemented a tool (Figure 1) to calculate an average agreement score per clustering approach by comparing a number of runs within each of the three clustering approaches (hierarchical^39^, k-means^40^, spectral clustering^41^) using multiple parameters (kernels, measures, algorithms) specifically based on the data set provided. The number of comparisons (*c*), between runs of the same approach, is an additional overarching parameter of this tool and contributes to the agreement score. For each comparison, the parameters of the two runs are drawn randomly from a predefined set (Table 1) selected randomly with replacement while not allowing the same parameters to be used within one comparison. In the interest of performance and computational time we suggest three comparisons to be used. Depending on *c*, we generate variations of the base clustering algorithms (package kernlab v0.9-29), along with the various distance measures and clustering categories they belong to. Within each pair of clustering runs the agreement is calculated using the adjusted Rand Index (package fossil v0.3.7), the corrected-for-chance version of the original Rand index^42^, which is based on the number of times any pair of points is partitioned in the same subgroup throughout different clusterings runs. To calculate the agreement within each clustering algorithm (spectral, k-means, hierarchical) we are considering pairs of runs using the same algorithm but different parameters. For those pairs the agreement is averaged across clustering runs and *k* number of clusters tested. The algorithm that presents the highest intra-method agreement over a logical range of clusters (*k* ∈ [2, *x*]) is noted as the most appropriate clustering of the samples based on a detected signal. A logical range of *k* is considered a set of successive *k’s* (where *k*⩾2) that is most probable to exist within our data, often determined by prior knowledge of the data, previous studies or domain expertise. This selection procedure is mainly affected by the type and size of the data leading similar datasets to opt for the same method due to the specific mathematical formulas within each algorithm.

**Table 1.** The clustering algorithms, their approach category and the various distance measures tested

| Clustering algorithms | Category | Distance measures/kernels | Additional parameters |
| --- | --- | --- | --- |
| K-means | Partitioning | Hartigan-Wong, Lloyd, Forgy, MacQueen | - |
| Hierarchical | Hierarchical | Euclidean, Manhattan, Minkowski, Canberra | Average, complete, median (linkage) |
| Spectral | Graph Theory | Rbfdot, Polydot, Tanhdot, Laplacedot, Vanilladot, Anovadot, Splinedot | - |

Spectral clustering algorithm^41^

Given a set of points S = {s_1_…,s_n_} in R^1^ that we want to cluster into **k** subsets:

1. *Form the affinity matrix A* ∈ *R^n**×**n^ defined by A_ij_ = exp(-||s_i_ - s_j_||^2^/2σ^2^) if i ≠ j, and A_ii_ = 0*
2. *Define D to be the diagonal matrix whose (i, i)-element is the sum of A’s i-th row, and construct the matrix L = D^-l/2^AD^-l/2^*
3. *Find x_1_, x_2_,…, x_k_, the k largest eigenvectors of L (chosen to be orthogonal to each other in the case of repeated eigenvalues), and form the matrix X = [x_1_ x_2_…x_k_] ∈ ^n**×**k^ by stacking the eigenvectors in columns*
4. *Form the matrix Y from X by renormalizing each of X’s rows to have unit length (i.e. Y_ij_ = X_ij_ / (Σ_j_ X^2^_ij_)^1/2^)*
5. *Treating each row of Y as a point in R^k^, cluster them into k clusters via K-means or any other algorithm (that attempts to minimize distortion)*
6. *Finally, assign the original point s_i_ to cluster j if and only if row i of the matrix Y was assigned to cluster j*

Hierarchical clustering algorithm (average linkage)

Given a set of points S = {s_1_…, s_n_} that we want to cluster into **k** subsets:

1. *Initialize with n clusters, each containing one data point (s_i_*)
2. *Compute the between-cluster distance D(r, s) as the between-object distance of the two data points in clusters r and s respectively, r, s =1, 2,…, n. Let the square matrix D = (D(r, s)). Various distances can be used (euclidean, manhattan, canberra, minkowski, maximum).*
3. *Find the most similar pair of clusters r and s, such that D(r, s) is minimum among all pairwise distances*
4. *Merge r and s to a new cluster t and compute the between-cluster distance D(t, k) for any existing cluster k ≠ r, s. Once the distances are obtained, delete the rows and columns corresponding to the old cluster r and s in the D matrix, as r and s do not exist anymore. Then add a new row and column in D corresponding to cluster t*.
5. *Repeat Step 3 a total of n - 1 times until there is only one cluster left*.
6. *Decide on a point to cut the cluster tree created above so as to obtain the desirable number of clusters (k)*

K-means^40^

K: kernel matrix, k: number of clusters, w: weights for each point, tmax: optional maximum number of iterations, {π_c_^(0)^}^k^_c=1_: optional initial clusters

1. *If no initial clustering is given, initialize the k clusters π_1_^(0)^,…, π_k_^(0)^ (i.e. randomly). Set t = 0*
2. *For each a_i_ and every cluster c, compute*

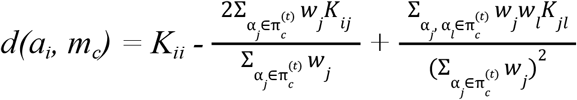
3. *Find c*(a) = argmin_c_d(a_i_, m_c_) resolving ties arbitrarily. Compute the updated clusters as*

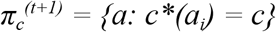
4. *If not converged or t_max_ > t, set t = t + 1 and go to Step 2; Otherwise, stop and output final clusters*

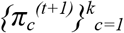

### Feature set subsampling

While gene expression data provide measures on the thousands of transcripts in the transcriptome, not all of them may provide discriminative information on the samples and may not be useful for clustering. Moreover, most clustering algorithms are heavily affected by a large number of features both computationally due to input size and in performance due to misdirecting data noise^43^. A common strategy to select interesting and potentially useful RNA features is to measure their variance across samples and select the ones with the highest scores instead of those that are either housekeeping or do not differentiate in our context. In this tool, we exclude RNA features that remain stable across samples and are therefore unable to offer any discriminatory power to our unsupervised machine learning models. Furthermore, the exhaustive feature selection procedure incrementally considers all the genes in the feature set and takes into account the stability of all generated test clusters and number of cluster ranges. This step does not require any deep knowledge or filtering decisions by the user.

Based on this observation our sample selection step, which is a part of the tool for bootstrap resampling of features presented in Figures 1C and 2A, first ranks features in a descending order of variance (var() function from the Stats R package) across samples, generating a list of the most variable features. Subsequently, multiple datasets of all samples and subsets of features are generated. All subsets draw a different number of features from the top of the variance list with replacement. The first dataset uses a relatively small number of features (*n*), depending on the total number of features (*N*) and the granularity of the result desired. The following datasets re-draw from the initial list increasing the number of features by *n*, ending up with 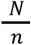 datasets.

**Figure 2:**
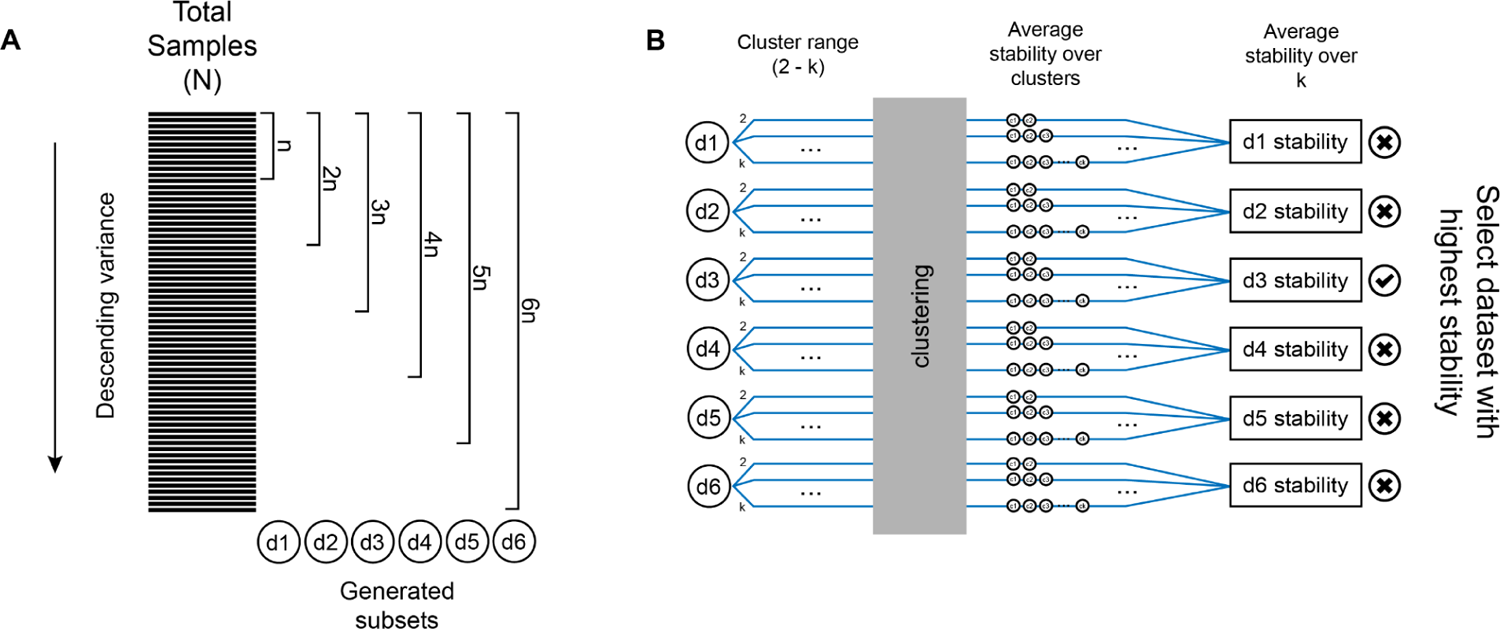
Sample selection overview. (A) Ranking of samples based on their variance across features and the subsequent generation of datasets of increasing size. (B) Calculation of the stability score of each generated dataset. Initially, we select a cluster range to run our clustering method for each dataset. After the clustering procedure, we calculate and average the stability over the generated clusters. Finally, we average the stabilities over k per dataset and determine a final stability score for each dataset. The features of the dataset with the highest stability are the ones that compose the most appropriate set for the downstream pipeline

### Stability-based assessment of feature sets

To assess the suitability of each resampled feature set for our clustering, we measure the average stability of the clusters they generate per run when a clustering method is applied over a range of *k*’s (Figure 2B). First, the clustering range, where the stability of each dataset will be calculated, is selected. For each dataset we generate the bootstrap stability for every *k* within range. To calculate each bootstrap stability score the data is randomly sampled with replacement and clustered internally using a spectral approach. We then compute the Jaccard similarities between the original clusters and the most similar clusters in the resampled data. The above procedure results in a stability score for each *k* and each dataset. We then calculate the final stability of each dataset by averaging the stability over *k*. The genes that comprise the dataset with the highest stability are the ones that compose the most appropriate set for the downstream analysis.

### Choosing k number of clusters

Most clustering methods require the number of *k* clusters to be defined as a parameter before the application of the algorithm on the data. The lack of a concrete way to determine the real number of clusters in a dataset led many studies to base their estimation on field/prior knowledge or various estimation methods such as the Silhouette score^44^. However, each method favours different aspects of the generated clusters (i.e. how compact clusters are and how far apart cluster centres are) and therefore suits specific datasets and may introduce bias towards the selection of *k*. To encompass these different angles in one methodology, avoid the risk of selecting an ineffective index and present a more general solution, this tool uses an ensemble learning approach (Figure 1) where multiple internal cluster indexes contribute to the decision making^45^. This approach prevents any bias from specific metrics and frees the user from making decisions on any specific metric and assumptions on the optimal number of clusters.

Initially, the value of the 15 indexes is calculated for each *k* within a cluster range of [2, *x*], where *x* is a logical upper limit of the number of clusters realistic for our dataset. The means over *k* are calculated per index and the optimal *k* is estimated by majority voting of the 14 means that evaluate the compactness and/or the distance between different subgroups. The selection of indexes can be found in Table 2. It is important to note that the most important aspect of determining *k* is minimum loss of information which directs us to overestimate and not underestimate *k^43^* while interpreting the voting results. Furthermore, cases that present only a single *k* as the optimal number of clusters should be treated with caution in case they are a result of a biased dataset.

**Table 2:** The list of 15 internal indexes used to estimate the optimal number of clusters (k).

| Internal index | Ideal | Formula | Source |
| --- | --- | --- | --- |
| Calinski-Harabasz | max | $\frac{\left(\frac{\text{between cluster variation}}{k-1}\right)}{\left(\frac{\text{within cluster variation}}{n-k}\right)}$ | 46 |
| Dunn | max | $\frac{\min(\text{intercluster distance})}{\max(\text{distance between all pairs})}$ | 47 |
| Pbm | max | $\frac{1}{k} * \frac{E^T}{E^W} * D_B$ | 48 |
| Tau | max | $\frac{\text{concordant pairs} - \text{discordant pairs}}{\sqrt{N_B * N_W \frac{N_T(N_T-1)}{2}}}$ | 49 |
| Gamma | max | $\frac{\text{concordant pairs} - \text{discordant pairs}}{\text{concordant pairs} + \text{discordant pairs}}$ | 50 |
| C index | min | $\frac{S_W - S_{MIN}}{S_{MAX} - S_{MIN}}$ | 51 |
| Davies–Bouldin | min | $\frac{1}{n} * \sum_{i=1}^n \max(\frac{\text{clusters scatter difference}}{\text{cluster separation}})$ | 52 |
| Mcclain<br>rao | min | $\frac{N_B}{N_W} * \frac{S_W}{S_B}$ | 53 |
| sd_dis | min | $\alpha * (\text{avg scattering for clusters}) + \text{total separation between clusters}$ | 54 |
| Ray–Turi | min | $\frac{1}{n} * \frac{\text{within-cluster dispersion}}{\text{min of the sq. distances between all the cluster barycenters}}$ | 55 |
| g_plus | min | $\frac{2 * s^-}{N_T(N_T - 1)}$ | 56 |
| Silhouette | max | $\text{average distance between clusters}$ | 44 |
| s_dbw | min | $\text{mean dispersion of clusters} + \text{between cluster density}$ | 57 |
| Compact<br>ness | max | $\text{Intra} - \text{Cluster distance}$ | 58 |
| Connecti<br>vity | max | $\text{The extent by which the items are placed in the same cluster as their nearest neighbours in the data space}$ | - |

All indexes are using different formulas to score a partitioning, measuring one or both of the following concepts: a) how compact each cluster is and b) how well the clusters separate. For each index we present which value is preferred (min or max) and its source. For the formulas: $k$ = number of clusters, $n$ = number of data points, $E^T$ = sum of the distances of all the points to the barycenter $G$ of the entire dataset, $E^W$ = sum of the distances of the points of each cluster to their barycenter, $N_B$ = pairs constituted of points which do not belong to the same cluster, $N_W$ = pairs constituted of points which belong to the same cluster, $N_T = N_W + N_B$ , $S_W$ = sum of the $N_W$ distances between all the pairs of points inside each cluster; $S_{MIN}$ = sum of the $N_W$ smallest distances between all the pairs of points in the entire data set, $S_{MAX}$ = sum of the $N_W$ largest distances between all the pairs of points in the entire data set, $S_B$ = sum of the between-cluster distances, $\alpha$ = weight equal to the value of average scattering of clusters obtained for the partition with the greatest number of clusters.

### Optimal parameter tuning

Previous steps have selected the optimal method, number of features and clusters. To perform the optimal clustering we automate the selection of parameters for each method so that manual tuning is not required. Towards that goal we utilise cluster stabilities to decide on the parameters (which depend on the specific algorithm i.e. kernels in k-means and spectral clustering, linkage method in hierarchical clustering) selected by this toolkit. All available parameters (Table 1) participate in the selection procedure where we measure the average bootstrap stability of the clusters (clusterboot function in R package fpc v2.2-3) using the previously determined optimal *k* and feature set for each parameter. The parameter that produces the highest stability is used for the optimal clustering run.

### Test datasets

Five datasets were used to validate different capabilities of the Omada package. First, two datasets were simulated by Omada’s functions. Function feasibilityAnalysisDataBased() was used to generate a multi-class dataset with 359 samples and 300 genes based on the contents and dimensions of the original RNA-seq data^18^ and composed of five groups of samples drawn from five different distributions with means (5,16,27,38,50) and sd (1,3,5,7,10), representing the five classes. Function feasibilityAnalysis() simulated a single-class dataset of 100 samples and 100 genes drawn from a single distribution. For the multissue Pan-cancer dataset we downloaded RNAseq expression data for 2244 samples and 253 genes representing three types of cancers: breast (n=1084), lung (n=566) and colon/rectal (n=594) downloaded through cbioportal^59^ from TCGA PanCancer Atlas^60^. The mRNA expression was in the form of z-scores relative to normal samples where we applied an extra step of arcsine normalisation. After filtering for tissue-specific genes^61^ for the three cancer types we retained 243 genes. Next, we utilised a PAH dataset (25,955 genes) generated from 359 patient samples with idiopathic and heritable pulmonary arterial hypertension (IPAH/HPAH). The transcriptomic data can be found in the EGA (the European Genome-phenome Archive) database under accession code EGAS0000100553265^62^ (restricted access) and all pre-processing details and parameters used can be found in ^18^. Finally, we used an RNA dataset from the whole blood of 238 mothers during midgestation (26-28 weeks of pregnancy). Read counts were extracted from GEO (accession number GSE182409^63^) and were then read into R and converted into TPM using the *convertCounts* function available in the *DGEobj.utils package*. For the purpose of clustering, we mapped the TPM dataset to the list of 24,070 genes used in the PAH dataset described in a previous section.

## Results

Omada was applied to five diverse gene expression datasets to demonstrate its utility in guiding cluster analysis and identifying plausible subgroups of samples. Two datasets were simulated by our tools. The simulated dataset with multiple distinct classes was used to determine Omada’s ability to accurately estimate *k* with reasonable stability when we know the existence of sample classes. In contrast, samples in the single-class simulated dataset were drawn from a single class and used to demonstrate the toolkit’s ability to point towards the lack of sample subgroups by indicating inconclusive low scores throughout the analysis. A multi-tissue pancan dataset was introduced to assess Omada’s capability to generate signal-based clusters that closely follow the tissue-specific patient sample distributions. In addition, to determine whether Omada can identify distinct heterogeneous subgroups from data without any prior classification information but potential present heterogeneity, we used a whole blood RNA-seq dataset from patients with pulmonary arterial hypertension (PAH)^18^. Lastly, implementation of the toolkit on a whole blood expression dataset (GUSTO) was included to demonstrate a case with potential technical biases and no known subgroups since it is composed of healthy participants.

For the above, we measured its consistency on algorithm, feature and number of clusters (*k*) selection and the stability of the generated clusters for a particular *k* (*stability_k_*) and the average across *k’s* (*stability_avg_*). It’s important to note that the value of this validation is derived from the fact that unstable clusters should not be interpreted as this instability comes from problematic data or an incorrect approach. However, cluster stability only provides a mechanistic way to assess the underlying data structure and further information is required to fully biologically validate the clusters^38^.

### Identifying known clusters

#### Multi-class dataset: Five distinct simulated expression classes representing heterogeneity

Omada’s basic function is to help identify samples that come from different sources and group together samples that come from the same source. Towards that end, we simulated a dataset with five sets of expression profiles with 50 samples each and 120 genes sourced from five unique distributions of expression data that represent heterogeneity within our samples. The means and standard deviations of each class are presented in Figure 3A, depicting the expression differences. Additionally, the empirical Cumulative Distribution Functions (ECDFs) of the five simulated classes (Figure 3B) as well as the high average Kolmogorov-Smirnov distances (D_avg_=88.3%, supplementary Table 1) show distinct differences between the distributions in respect to the expression in the simulated RNA-seq dataset. To demonstrate the effect of different sample and gene numbers, multiple datasets were simulated with an increasing number of samples and genes (Figure 3C). The calculated cluster stabilities, where each value represents the stability over a range of *k* and a specific number of samples and features, show five or less samples per class provide highly unstable and unreliable clusters. The minimum acceptable stability threshold of 60% was achieved with at least 20 samples and a reliable stability of 75% was achieved using 1000 samples.

**Figure 3:**
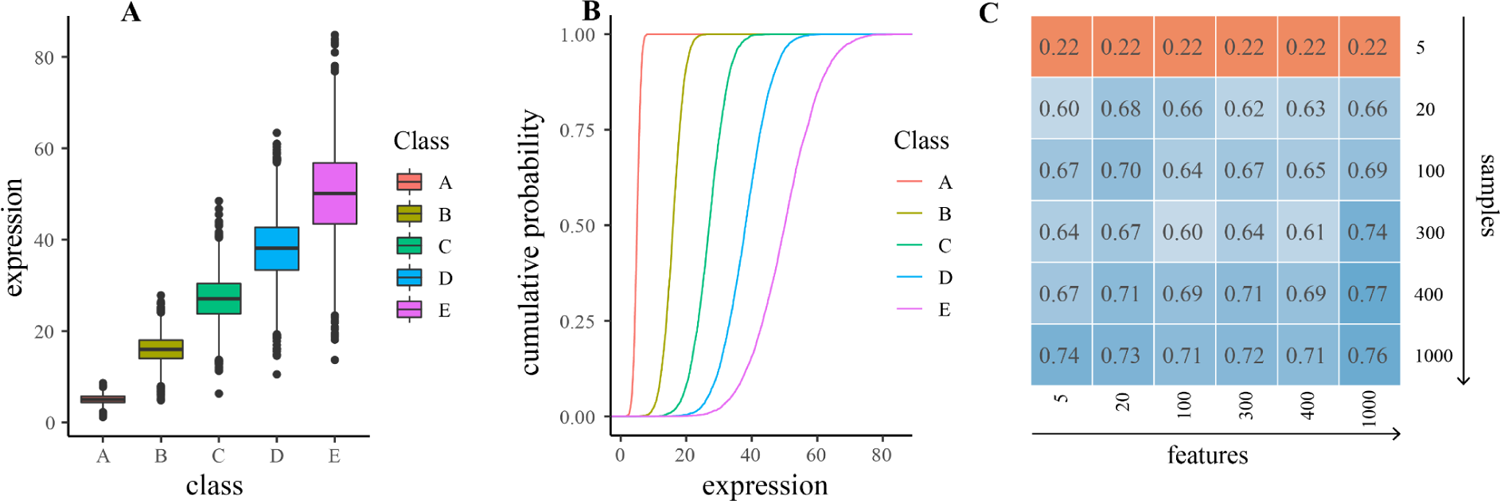
A) Expression boxplots for the five clusters showing the means and standard deviations B) The cumulative probability (as calculated from the empirical cumulative distribution function) for the five clusters calculated by a two sided Kolmogorov-Smirnov Test C) Average over-k stabilities for simulated datasets of increasing sample and gene numbers. A small number of samples consistently provides extremely unstable clusters (orange) while increasing both numbers consistently produces datasets that pass the stability threshold of 0.6 (blue).

To test the ability of the clustering tools to produce stable clusters in various contexts we first apply them in sequence on strategically simulated data. The data are composed of distinct classes (based on class mean *m_class_* and standard deviation *sd_class_*) and due to that strong signal our tools are expected to determine an accurate *k* with reasonable stability, scoring above 60%. To allow for a more direct comparison, we used a multi-class simulated dataset (see *Test datasets* in *Methods*) based on the original RNA-seq data^18^. When considering ranges of *k* we are using [2, 6] clusters to observe a broader range of results for comparison reasons. First, the clustering feasibility tool showed that the highest stability was 78% (Supplementary Table 4) providing a strong indication of stability across our clusters. Since we selected a limited range of *k* ∈ [2, 6] where the stability should remain high, the averaged -over every tested *k*- stability (stability_avg_) of 72% indicates a dataset of adequate size and class definition to proceed to clustering analysis. It should be noted that when large ranges of *k* are selected the average stability will naturally decrease as the calculations will take into account *k*’s much larger or smaller than the actual number of classes in the data. In such cases the user can review the individual *k* stabilities generated as part of this tool to conclude whether those values are satisfying i.e a minimum of 60%. Next, we calculated the partitioning agreement of three clustering algorithms and spectral clustering showed the highest average score of 56% (Figure 5A and Supplementary Table 4). Partitioning agreement scores should be interpreted across algorithms applied on the same dataset rather than as absolute values keeping in mind that a score below 50% represents a random partitioning and subsequently a non-robust clustering. In the subsequent feature selection step, the highest average stability was registered when using all 300 features (stability_avg_ = 78%, Supplementary Table 4), not discarding any feature as they all demonstrated very similar variance due to the nature of the simulated data. Finally, 8 out of 15 internal metrics voted five clusters as the optimal number during the k estimation step (Figure 5B) providing a confident estimation above 50%.

**Figure 4:**
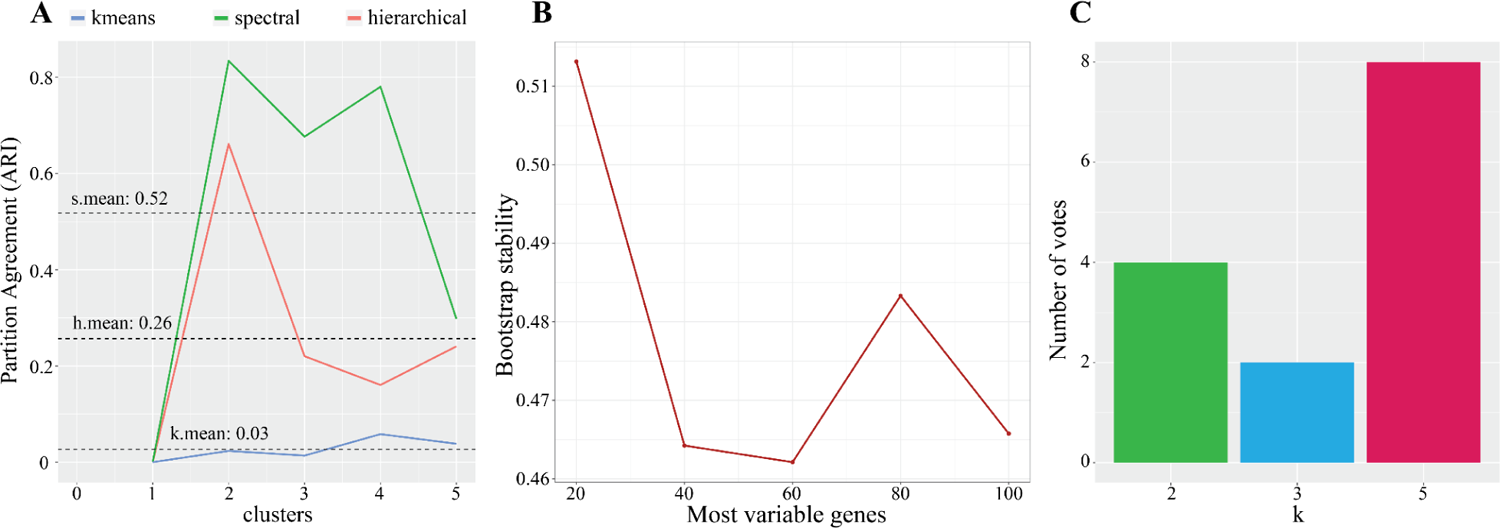
Performance criteria for single-class simulated dataset. The results demonstrate low scores for the majority of steps. A) shows the average partition agreement of all three algorithms below the 52% mark indicating very unstable clustering runs overall. In B) the stability of every possible subsets of genes does not surpass 51.3% underlying overall unstable clusters. C) shows five clusters as the first estimate (voted by 8 metrics), significantly different from the one class this dataset contains.

**Figure 5:**
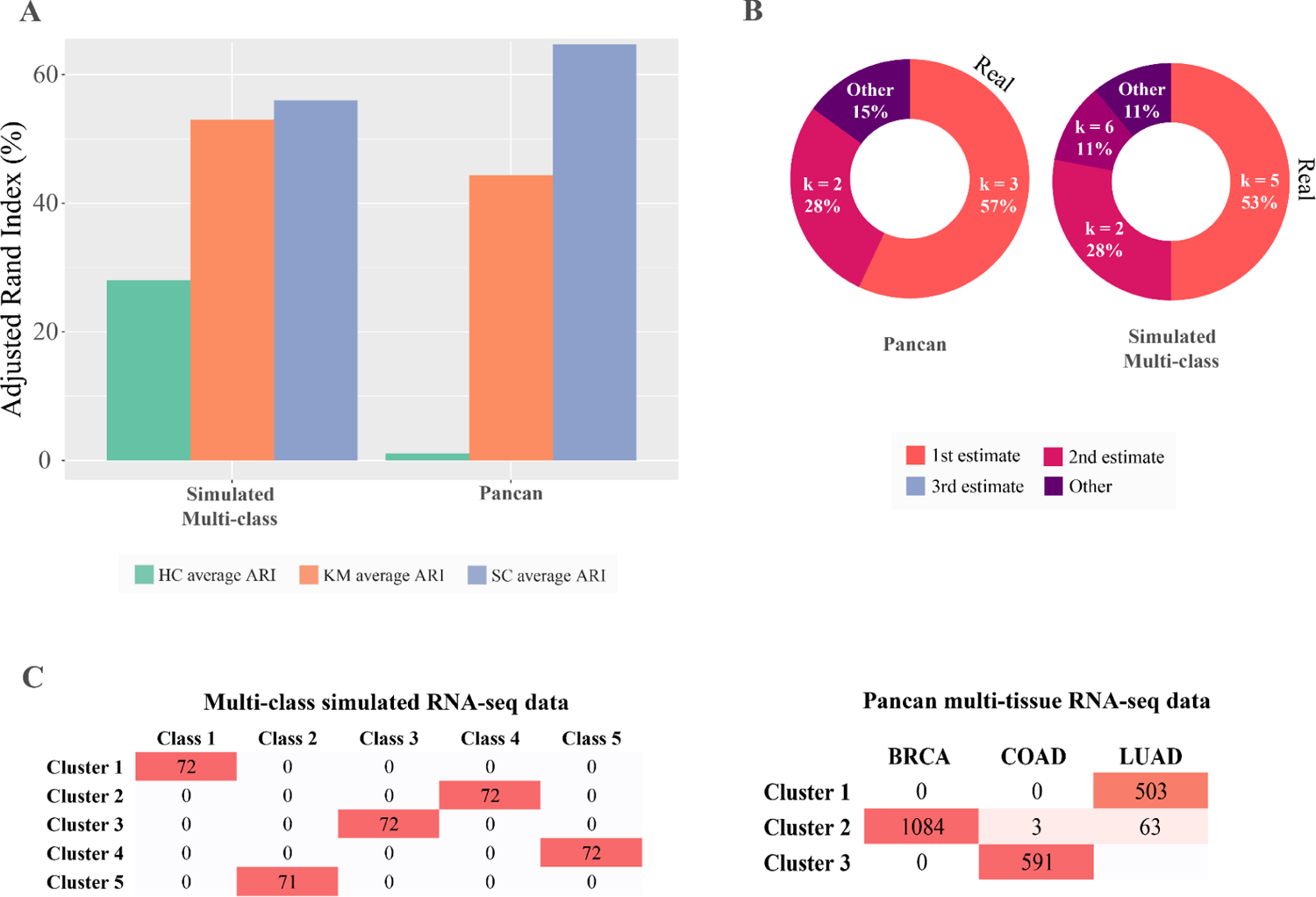
Performance criteria for two heterogeneous datasets, simulated multi-class and Pancan dataset. The multi-class dataset contains artificial samples from five distinct clusters and the Pancan dataset is composed of three different cancer types presenting biological heterogeneity. A) The agreement between the predicted and true clusters (Adjusted Rand Index) from three different clustering algorithms (HC: hierarchical, KM: K-means, SC: spectral clustering) applied to the two datasets. B) Shows the real number of clusters for the dataset (black text) and the three most likely number of clusters k, with estimates of their percent probability. C) The contingency tables of the combinations between generated clusters (1st estimates) and real classes in the datasets. Darker red colour intensity denotes higher frequency.

#### Single-class dataset: Homogeneously simulated dataset

To demonstrate Omada’s ability to identify datasets without any present clusters where all patients belong to one class, we used the single-class simulated dataset (see *Test datasets* in *Methods*). All potential *k* of two or higher achieved low scores with average and maximum stabilities of 45% and 55%, respectively (supplementary Table 2). It is recommended to avoid clustering analysis on such low score datasets and instead opt for scores of at least 60%. Ideally, stabilities of 80-90% are considered very strong^64^, however the potential of several signals in transcriptomic data and the exploration across multiple *k* generally decreases the output stability to an acceptable threshold of 60-70%. Next, Figure 4A shows the overall low partitional consistencies (averaged over all tested *k*) for all algorithms with spectral average partition agreement of 52%, kmeans average partition agreement of 3% and hierarchical average partition agreement of 26%. With the best performing algorithm showing an agreement of around 50% we can assume that the tested algorithms are randomly assigning memberships, therefore we cannot achieve a robust model with the current data. When using spectral clustering to select the most appropriate set of genes, the cluster stability rapidly dropped below 50% when using more than 20 genes (Figure 4B) indicating that the algorithm got worse in assigning memberships as we considered more simulated genes. Finally, the ensemble voting step showed the majority of the votes supporting five clusters (Figure 4C), a significant variation from the single simulated class of this dataset. In such unexpected outputs, one should examine the generated metric scores. In this case, the vast majority of metric scores are worse when we are testing single-class instead of multi-class simulations (supplementary Table 3) inferring lower cluster quality, i.e lower compactness and smaller distance between clusters. Additionally, worse scores (decided according to Table 2) infer higher uncertainty during the voting process.

### Discriminating cancer types from pan cancer tissue expression data

An integral capability of Omada is to accurately stratify patients according to any biologically relevant signal present in expression data and detect differences stemming from genes, pathways, tissues etc. Real multi-tissue samples are often the focus of exploratory studies as they present cell-type differences but still unknown factors that may discriminate them. Using expression data from multiple cancer types (Pan-cancer dataset as described in *Test datasets* in *Methods*), we expect our tools to identify clusters that are consistent with the samples’ tissues of origin. Due to the different types of tumours we explored the potential cluster range of [2, 5] for each pipeline step. The clustering feasibility of the dataset (2244 samples, 243 genes) presented an average stability of 88% and maximum stability of 100% (Supplementary Table 5) providing confidence for the downstream analysis. Spectral clustering showed the highest consistency (partition agreement_avg_ = 63% closely resembling the simulated multiclass dataset, Figure 5A) and was therefore deemed as the most robust. In this example hierarchical clustering showed high instability, as shown in Figure 5A, demonstrating the importance of selecting the appropriate algorithm to create a robust model. According to our selection tool, all 243 genes produced the most stable set of clusters with a stability of 96% (Supplementary Table 5) which coupled with the high algorithm robustness indicated a model that most likely detects a signal in the data. Additionally, a very important observation is that all genes were deemed important to produce nearly perfectly stable clusters agreeing with the filtering of genes based on the cancer type annotations we performed prior to this clustering analysis. The ensemble voting tool estimated our dataset to contain three clusters of samples with the support of 57% of the metrics (Figure 5B). When comparing these results with the simulated five-class dataset, both achieved higher certainty on the five clusters (>50%, Figure 5B) reflecting the rigid differences between the clusters when dealing with cancer tissues and simulated classes. In the case of the pancan partitioning, the breast, lung and colon/rectal samples almost perfectly grouped in their respective clusters (Figure 5C).

### RNAseq data from diseased tissue with unknown heterogeneity

It is important for Omada to be able to robustly identify patient subgroups when heterogeneity for the cohort has not been previously characterised. We applied our tools on such a dataset (PAH dataset as described in *Test datasets* in *Methods*) to assess whether they can still produce stable clusters that differ in terms of expression profiles and other phenotypic measures. The feasibility for this dataset’s simulation showed an average stability of 61% and a maximum stability of 74% both acceptable to proceed with the clustering analysis (Supplementary Table 7). A notable 13% difference between average and maximum stability provides a positive indication that a specific *k* might prove significantly more stable downstream. The spectral clustering technique recorded the highest partitional consistency (partition agreement_avg_ = 86% and partition agreement_max_ = 96%, Supplementary Table 7) when we examined each algorithm’s partition agreement for up to ten clusters. The bootstrapping subset selection tool estimated the 300 most variable genes as the most stable clustering parameter with a maximum stability of 73% (Figure 6A) showing an impressive reduction from the initial gene set (25,955) and ensuring the removal of a lot of data noise. According to the ensemble voting tool two clusters were voted by 71% of the internal metrics followed by *k* = 3 (14%) and *k* = 5 (7%). Despite the strong indication of two clusters, *k* = 5 was selected to prevent loss of information occuring when smaller embedded clusters are disregarded. As shown by the downstream analysis, fully presented in ^18^, selecting the higher *k*, even as a second estimate, allowed us to detect strong expression profiles. After considering cluster sizes the three predominant subgroups showed significant differences in expression, immunity and survival profiles as well as risk category distributions (Figure 6B, C).

**Figure 6:**
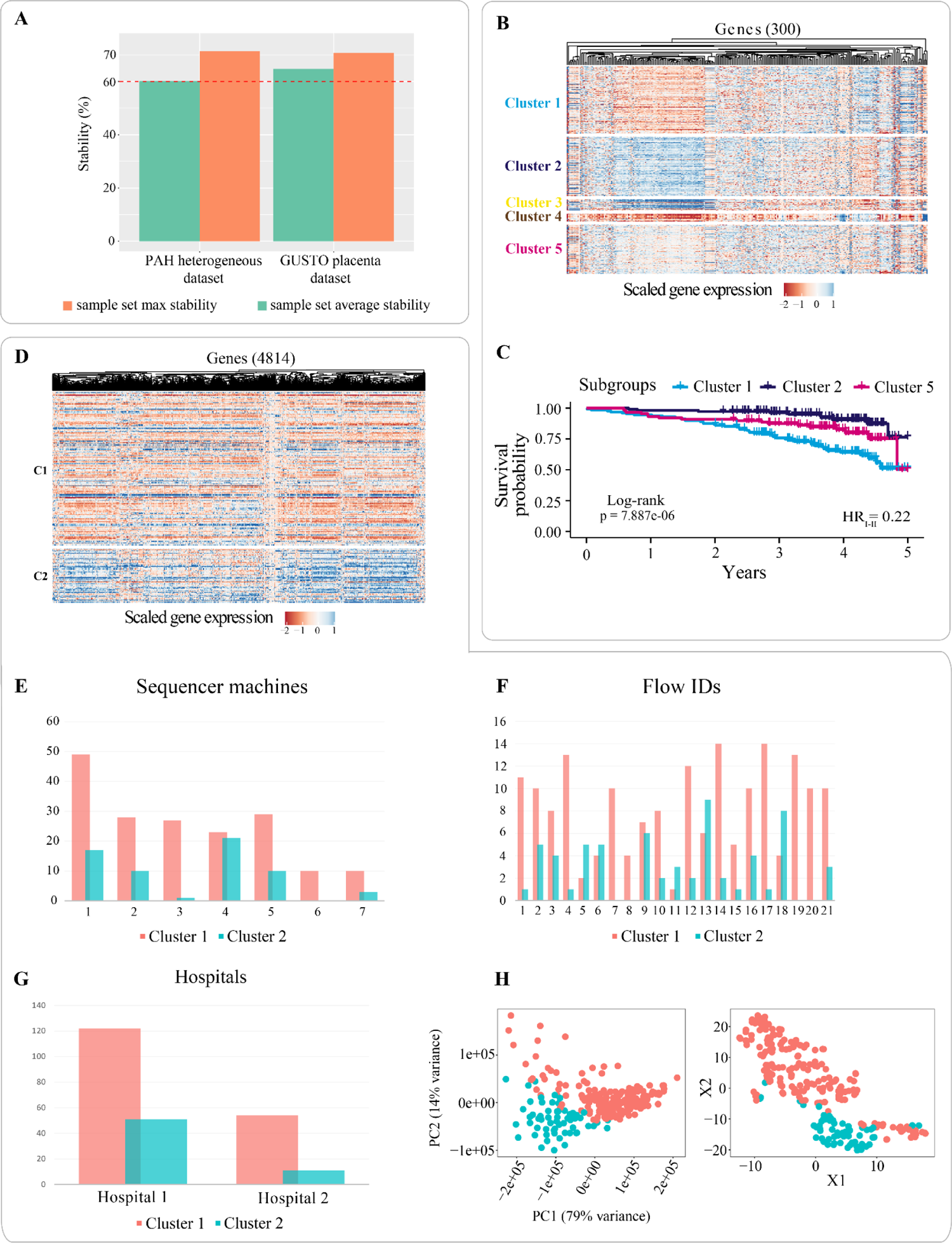
Performance criteria for PAH and GUSTO datasets which have no known subgroups. Panel A) depicts the average and max sample set stabilities (percentage) for both datasets. The red dashed line represents the threshold of a stable clustering (60%). The PAH RNA-seq dataset contains expression of IPAH patients with panel B) showing the gene expression heatmap and C) survival profiles for discovered subgroups. The GUSTO dataset contains expression from healthy maternal whole blood with panel D) showing the gene expression heatmap. The following panels show the distribution of cluster members across E) sequencer machines(chi-square p-value 1.39e-03), F) flow IDs (chi-square p-value 2.55e-06) and G) hospitals where the data were collected (chi-square p-value 0.072). H) t-SNE and PCA plots of the expression profiles with labelling of the two discovered subgroups.

### RNA-seq data from healthy whole blood tissue

Next we tested how Omada would discriminate samples from healthy individuals from a single tissue type. Generally, in studies based on a dataset with no discernible heterogeneity to be explored - i.e a dataset without patients of dissimilar outcomes or controls - clustering algorithms may not be robust and may generate variable results. Useful partitionings might still be formed, such as unforeseen disease subgroups, but these observations must be validated. Towards that end we used the GUSTO RNA dataset of 238 mothers, as seen in *Test datasets* in *Methods*. During determining clustering potential our simulated dataset showed stability_avg_ = 56% and stability_max_ = 59% (Supplementary Table 8), a similar low-stability score as in the simulated single-class (45% and 55%). We examined a k-range of [2, 5] where spectral and k-means clustering showed very similar internal average partitional agreements of 61% and 60% and very high maximum agreements of 93% and 88% (Supplementary Table 8), respectively. The extremely high agreement scores should be interpreted with caution as they might not reflect a very strong signal but an underlying bias that partitions samples in similar groups repeatedly, over-powering the parameter changes. The 50 most variable genes were estimated to produce the most stable clustering with maximum stability = 71% (Figure 6A). Similarly to the agreement scores, a small number of genes driving the most stable clusters (starting from 24,070 genes) might indicate either a strong expression signal or a pre-existing bias. When estimating the number of clusters, two (46%) and three (40%) clusters were voted by the majority showing a general consensus. Considering all the above strong indications, we need to assess the dataset and the resulting subgroups for potential biases before relying on the cluster memberships. Towards that end and utilising clinical data, the association results show the dataset might be biased based on technical batches with sequencer machine and flowID presenting significant differences between clusters (1.39e-03 and 2.55e-06, respectively) with hospital location coming close to significance with p-value = 0.072 (Supplementary Table 9). Additional statistical tests and regression analysis with maternal and foetal physiological and clinical phenotypes did not show any association with the clusters. The expression profiles of the two clusters show visible differences as do the t-SNE and PCA analyses (Figure 6E) with the first principal component of the latter explaining 79% of the variance in the GUSTO dataset.

## Discussion

Our toolkit is designed to answer multiple questions arising during transcriptomic exploratory studies that target to uncover heterogeneity and subtypes within any condition that might be driven by expression changes. With the plurality of unsupervised methods available and their specialised nature, the selection of the most appropriate approach is a multi-factor problem. A lot of technical decisions are required in the procedure starting with a dataset and completed with a meaningful set of subgroups. To assist with this problem, our toolkit initially assesses the potential of a target dataset and provides estimates of the most appropriate method, gene set and number of subgroups finally outputting a partition based on optimised parameters unburdening the user of specialised decision making (Figure 1). All individual tools are computed internally and therefore do not require prior deep knowledge of machine learning by the user. All results, intermediate and final, are observable and each step is justified by multiple measures and indices representing widely used machine learning techniques.

Applying unsupervised learning on expression datasets is often not a straightforward task as it contains the element of uncertainty mainly introduced by the lack of knowledge on the data points. No methods or metrics can give a definitive answer to the main clustering questions, as presented in previous sections, therefore each tool has to be used with caution, i.e. determining the dataset clustering potential is an indication rather than a clear sign that partitioning the dataset will yield informational subgroups. Additionally, clustering can often contain non-deterministic steps allowing for each function to behave slightly differently between similar runs. To reduce the uncertainty and provide a reliable set of tools, this toolkit has been applied on various gene expression datasets where its efficiency has been demonstrated. However, it is important to note that despite the use of multiple methodological approaches within this toolkit the inherent exploratory characteristics of clustering do not allow for clusters of definite value, instead they are meant to be dealt with scientific caution and biological validation. Aside from the actual memberships, the functions in this package can also reveal useful information about the input data. The GUSTO RNA-seq dataset showed that biases can be discovered by applying simple tests, such as PCA or tSNE, in conjunction with the cluster members. It is also possible for Omada to hint towards the existence of a single class, and therefore no heterogeneity, by consistently revealing low partition agreement and stability scores across multiple functions, as demonstrated in our single-class dataset example. Furthermore, Omada can help in selecting a small group of genes with potential partitioning capabilities as the feature selection step is expected to greatly reduce the number of genes which in most cases count to thousands. This toolkit is currently based on specific clustering techniques and metrics but its modular nature allows its extension to accommodate different data types that come in the form of continuous numeric values such as microRNA, metabolite or single cell RNA datasets. Furthermore, the structure of this toolkit allows for additional approaches to be added in the future to the pool of clustering algorithms to be tested keeping up with the current state of the art techniques.

## Code Availability

Code will be available on github and as a bioconductor software package (Omada) at 10.18129/B9.bioc.omada.

## Data Availability

The expression datasets used in this work can be accessed through the following sources: The two simulated, by Omada, datasets (single and multi-class) can be accessed and downloaded at https://github.com/BioSok/OmadaSimulatedDatasets. The Pan cancer tissue expression data can be accessed through (https://archive.ics.uci.edu/ml/datasets/gene+expression+cancer+RNA-Seq). The transcriptomic data used in this study can be accessed through the EGA (the European Genome-phenome Archive) database under accession code EGAS00001005532. In compliance with the Ethics under which these data and samples have been collected, the transcriptomic data are available through restricted access for approved researchers who agree to the conditions of use, i.e. keeping it secure and only using it for approved purposes. To apply for access please contact. You will receive an application form within 30 days. The ‘UK National PAH Cohort Study Data Access Committee’ will review requests within 3 months of receipt of the completed application form and if approved, provide details for access to the RNAseq data stored at the EGA. All requesters must agree to the data access conditions found in EGA. The data used to generate statistics, plots and figures are accessible through our interactive portal found in https://sheffield-university.shinyapps.io/ipah-rnaseq-app/. The GUSTO expression dataset is available in NCBI Gene Expression Omnibus (GEO; https://www.ncbi.nlm.nih.gov/geo/) under the accession numbers GSE182409 (Corresponding Reviewer token number: qjolmmeudnofnsv).

## Acknowledgements

The UK National Cohort of Idiopathic and Heritable PAH is supported by grants from the British Heart Foundation (SP/12/12/29836 & SP/18/10/33975) and the UK Medical Research Council (MR/K020919/1). Additional samples from the Sheffield Teaching Hospitals Observational Study of Pulmonary Hypertension, Cardiovascular and other Respiratory Diseases were supported by British Heart Foundation (PG/11/116/29288). S.K. is supported by a Donald Heath Ph.D. Studentship award and A*STAR Research Attachment Programme (ARAP) award.

The GUSTO study group includes Allan Sheppard, Amutha Chinnadurai, Anne Eng Neo Goh, Anne Rifkin-Graboi, Anqi Qiu, Arijit Biswas, Bee Wah Lee, Birit Froukje Philipp Broekman, Boon Long Quah, Chai Kiat Chng, Cheryl Shufen Ngo, Choon Looi Bong, Christiani Jeyakumar Henry, Daniel Yam Thiam Goh, Doris Ngiuk Lan Loh, Fabian Kok Peng Yap, George Seow Heong Yeo, Helen Yu Chen, Hugo P. S. van Bever, Iliana Magiati, Inez Bik Yun Wong, Ivy Yee-Man Lau, Jeevesh Kapur, Jenny L. Richmond, Jerry Kok Yen Chan, Joanna Dawn Holbrook, Johan G. Eriksson, Joshua J. Gooley, Keith M. Godfrey, Kenneth Yung Chiang Kwek, Kok Hian Tan, Krishnamoorthy Naiduvaje, Leher Singh, Lin Lin Su, Lourdes Mary Daniel, Lynette Pei-Chi Shek, Marielle V. Fortier, Mark Hanson, Mary Foong-Fong Chong, Mary Rauff, Mei Chien Chua, Michael J. Meaney, Mya Thway Tint, Neerja Karnani, Ngee Lek, Oon Hoe Teoh, P. C. Wong, Peter David Gluckman, Pratibha Keshav Agarwal, Rob Martinus van Dam, Salome A. Rebello, Seang Mei Saw, Shang Chee Chong, Shirong Cai, Shu-E Soh, Sok Bee Lim, Stephen Chin-Ying Hsu, Victor Samuel Rajadurai, Walter Stunkel, Wee Meng Han, Wei Wei Pang, Yap Seng Chong, Yin Bun Cheung, Yiong Huak Chan and Yung Seng Lee.

## Author Contributions

SK and DW conceived the tools. SK undertook computational work and drafted the work with DW. AL, CR, TPF and MW participated in the data acquisition of the work. All authors revised it critically for important intellectual content; and gave final approval of the version submitted for publication.

## Notes

### Competing Interest Statement

The authors have declared no competing interest.

### Summary of Updates

Include the toolkit's name to associate it with the relevant Bioconductor package.

## References

1. Yu, N. Y.-L. et al. Complementing tissue characterization by integrating transcriptome profiling from the Human Protein Atlas and from the FANTOM5 consortium. Nucleic Acids Res. 43, 6787–6798 (2015).

2. Keen, J. C. & Moore, H. M. The Genotype-Tissue Expression (GTEx) Project: Linking Clinical Data with Molecular Analysis to Advance Personalized Medicine. J Pers Med 5, 22–29 (2015).

3. Uhlén, M. et al. Proteomics. Tissue-based map of the human proteome. Science 347, 1260419 (2015).

4. Wang, L. et al. RNA sequencing-based longitudinal transcriptomic profiling gives novel insights into the disease mechanism of generalized pustular psoriasis. BMC Med. Genomics 11, 52 (2018).

5. Neff, R. A. et al. Molecular subtyping of Alzheimer’s disease using RNA sequencing data reveals novel mechanisms and targets. Sci Adv 7, (2021).

6. Saeidian, A. H., Youssefian, L., Vahidnezhad, H. & Uitto, J. Research Techniques Made Simple: Whole-Transcriptome Sequencing by RNA-Seq for Diagnosis of Monogenic Disorders. J. Invest. Dermatol. 140, 1117–1126.e1 (2020).

7. Tran, H. T. N. et al. A benchmark of batch-effect correction methods for single-cell RNA sequencing data. Genome Biol. 21, 12 (2020).

8. Xing, Q. R. et al. Unraveling Heterogeneity in Transcriptome and Its Regulation Through Single-Cell Multi-Omics Technologies. Front. Genet. 11, 662 (2020).

9. Firth, A. L., Mandel, J. & Yuan, J. X.-J. Idiopathic pulmonary arterial hypertension. Dis. Model. Mech. 3, 268–273 (2010).

10. Koirala, B. et al. Heterogeneity of Cardiovascular Disease Risk Factors Among Asian Immigrants: Insights From the 2010 to 2018 National Health Interview Survey. J. Am. Heart Assoc. 10, e020408 (2021).

11. Rivera-Andrade, A. & Luna, M. A. Trends and heterogeneity of cardiovascular disease and risk factors across Latin American and Caribbean countries. Prog. Cardiovasc. Dis. 57, 276–285 (2014).

12. Manchia, M., Cullis, J., Turecki, G., Rouleau, G. A. & Uher, R. The impact of phenotypic and genetic heterogeneity on results of genome wide association studies of complex diseases. PLoS One (2013).

13. Vidman, L., Källberg, D. & Rydén, P. Cluster analysis on high dimensional RNA-seq data with applications to cancer research - An evaluation study. PLoS One 14, e0219102 (2019).

14. Ren, Z., Wang, W. & Li, J. Identifying molecular subtypes in human colon cancer using gene expression and DNA methylation microarray data. Int. J. Oncol. 48, 690–702 (2016).

15. Sotiriou, C., Neo, S. Y. & McShane, L. M. Breast cancer classification and prognosis based on gene expression profiles from a population-based study. Proceedings of the (2003).

16. Lapointe, J. et al. Gene expression profiling identifies clinically relevant subtypes of prostate cancer. Proc. Natl. Acad. Sci. U. S. A. 101, 811–816 (2004).

17. Wu, F. et al. Single-cell profiling of tumor heterogeneity and the microenvironment in advanced non-small cell lung cancer. Nat. Commun. 12, 2540 (2021).

18. Kariotis, S. et al. Biological heterogeneity in idiopathic pulmonary arterial hypertension identified through unsupervised transcriptomic profiling of whole blood. Nat. Commun. 12, 7104 (2021).

19. Xu, D. & Tian, Y. A Comprehensive Survey of Clustering Algorithms. Annals of Data Science vol. 2 165–193 Preprint at https://doi.org/10.1007/s40745-015-0040-1 (2015).

20. Reddy, C. K. & Vinzamuri, B. A Survey of Partitional and Hierarchical Clustering Algorithms. Data Clustering 87–110 Preprint at https://doi.org/10.1201/9781315373515-4 (2018).

21. Jamail, I. & Moussa, A. Current State-of-the-Art of Clustering Methods for Gene Expression Data with RNA-Seq. in Applications of Pattern Recognition (IntechOpen, 2020).

22. Ezugwu, A. E. et al. A comprehensive survey of clustering algorithms: State-of-the-art machine learning applications, taxonomy, challenges, and future research prospects. Eng. Appl. Artif. Intell. 110, 104743 (2022).

23. Cao, Y., Geddes, T. A., Yang, J. Y. H. & Yang, P. Ensemble deep learning in bioinformatics. Nature Machine Intelligence 2, 500–508 (2020).

24. Park, C., Took, C. C. & Seong, J.-K. Machine learning in biomedical engineering. Biomed Eng Lett 8, 1–3 (2018).

25. Choy, G. et al. Current Applications and Future Impact of Machine Learning in Radiology. Radiology 288, 318–328 (2018).

26. Stafford, I. S. et al. A systematic review of the applications of artificial intelligence and machine learning in autoimmune diseases. NPJ Digit Med 3, 30 (2020).

27. Hulsen, T. et al. From Big Data to Precision Medicine. Front. Med. 6, 34 (2019).

28. Wang, X., Williams, C., Liu, Z. H. & Croghan, J. Big data management challenges in health research—a literature review. Brief. Bioinform. 20, 156–167 (2019).

29. Nayyar, A., Gadhavi, L. & Zaman, N. Machine learning in healthcare: review, opportunities and challenges. Machine Learning and the Internet of Medical Things in Healthcare 23–45 Preprint at https://doi.org/10.1016/b978-0-12-821229-5.00011-2 (2021).

30. Gaba, D. & Mittal, N. 2. Implementation and classification of machine learning algorithms in healthcare informatics: approaches, challenges, and future scope. Computational Intelligence for Machine Learning and Healthcare Informatics 21–34 Preprint at https://doi.org/10.1515/9783110648195-002 (2020).

31. Wang, C., Gao, X. & Liu, J. Impact of data preprocessing on cell-type clustering based on single-cell RNA-seq data. BMC Bioinformatics 21, 440 (2020).

32. Eijssen, L. M. T. et al. User-friendly solutions for microarray quality control and pre-processing on ArrayAnalysis.org. Nucleic Acids Res. 41, W71–6 (2013).

33. Andrews, S. & Others. FastQC: a quality control tool for high throughput sequence data. Preprint at (2010).

34. Bolger, A. M., Lohse, M. & Usadel, B. Trimmomatic: a flexible trimmer for Illumina sequence data. Bioinformatics vol. 30 2114–2120 Preprint at https://doi.org/10.1093/bioinformatics/btu170 (2014).

35. D’haene & Hellemans. The importance of quality control during qPCR data analysis. Int. Drug Discov.

36. Baccarella, A., Williams, C. R., Parrish, J. Z. & Kim, C. C. Empirical assessment of the impact of sample number and read depth on RNA-Seq analysis workflow performance. BMC Bioinformatics 19, 423 (2018).

37. Wang, J., Yue, S., Yu, X. & Wang, Y. An efficient data reduction method and its application to cluster analysis. Neurocomputing 238, 234–244 (2017).

38. Hennig, C. Cluster-wise assessment of cluster stability. Comput. Stat. Data Anal. 52, 258–271 (2007).

39. Hartigan, J. A. Clustering Algorithms. (John Wiley & Sons, Inc., 1975).

40. Dhillon, I. S. A Unified View of Kernel K-means, Spectral Clustering and Graph Cuts. (Computer Science Department, University of Texas at Austin, 2004).

41. Ng, A. Y., Jordan, M. I. & Weiss, Y. On Spectral Clustering: Analysis and an algorithm. in Advances in Neural Information Processing Systems 14 (eds. Dietterich, T. G., Becker, S. & Ghahramani, Z.) 849–856 (MIT Press, 2002).

42. Rand, W. M. Objective Criteria for the Evaluation of Clustering Methods. J. Am. Stat. Assoc. 66, 846–850 (1971).

43. Rodriguez, M. Z. et al. Clustering algorithms: A comparative approach. PLoS One 14, e0210236 (2019).

44. Rousseeuw, P. J. Silhouettes: A graphical aid to the interpretation and validation of cluster analysis. Journal of Computational and Applied Mathematics vol. 20 53–65 Preprint at https://doi.org/10.1016/0377-0427(87)90125-7 (1987).

45. Polikar, R. Ensemble based systems in decision making. IEEE Circuits and Systems Magazine 6, 21–45 (2006).

46. Calinski, T. & Harabasz, J. A dendrite method for cluster analysis. Communications in Statistics - Theory and Methods vol. 3 1–27 Preprint at https://doi.org/10.1080/03610927408827101 (1974).

47. Dunn†, J. C. Well-Separated Clusters and Optimal Fuzzy Partitions. Journal of Cybernetics vol. 4 95–104 Preprint at https://doi.org/10.1080/01969727408546059 (1974).

48. Pakhira, M. K., Bandyopadhyay, S. & Maulik, U. Validity index for crisp and fuzzy clusters. Pattern Recognition vol. 37 487–501 Preprint at https://doi.org/10.1016/j.patcog.2003.06.005 (2004).

49. Kendall, M. G. A New Measure of Rank Correlation. Biometrika vol. 30 81 Preprint at https://doi.org/10.2307/2332226 (1938).

50. Baker, F. B. & Hubert, L. J. Measuring the Power of Hierarchical Cluster Analysis. Journal of the American Statistical Association vol. 70 31–38 Preprint at https://doi.org/10.1080/01621459.1975.10480256 (1975).

51. Hubert, L. J. & Levin, J. R. A general statistical framework for assessing categorical clustering in free recall. Psychological Bulletin vol. 83 1072–1080 Preprint at https://doi.org/10.1037//0033-2909.83.6.1072 (1976).

52. Davies, D. L. & Bouldin, D. W. A Cluster Separation Measure. IEEE Transactions on Pattern Analysis and Machine Intelligence vol. PAMI-1 224–227 Preprint at https://doi.org/10.1109/tpami.1979.4766909 (1979).

53. McClain, J. O. & Rao, V. R. CLUSTISZ: A program to test for the quality of clustering of a set of objects. J. Mark. Res. 12, 456–460 (1975).

54. Halkidi, M., Batistakis, Y. & Vazirgiannis, M. On Clustering Validation Techniques. J. Intell. Inf. Syst. 17, 107–145 (2001).

55. Ray, S. & Turi, R. H. Determination of number of clusters in k-means clustering and application in colour image segmentation. in Proceedings of the 4th international conference on advances in pattern recognition and digital techniques 137–143 (Citeseer, 1999).

56. Rohlf, F. J. Methods of Comparing Classifications. Annual Review of Ecology and Systematics vol. 5 101–113 Preprint at https://doi.org/10.1146/annurev.es.05.110174.000533 (1974).

57. Halkidi, M. & Vazirgiannis, M. Clustering validity assessment: finding the optimal partitioning of a data set. Proceedings 2001 IEEE International Conference on Data Mining Preprint at https://doi.org/10.1109/icdm.2001.989517.

58. Song, Y. Class compactness for data clustering. in 2010 IEEE International Conference on Information Reuse & Integration 86–91 (IEEE, 2010).

59. cBioPortal for Cancer Genomics. https://www.cbioportal.org/datasets.

60. Weinstein, J. N. et al. The Cancer Genome Atlas Pan-Cancer analysis project. Nature Genetics vol. 45 1113–1120 Preprint at https://doi.org/10.1038/ng.2764 (2013).

61. Kim, P. et al. TissGDB: tissue-specific gene database in cancer. Nucleic Acids Res. 46, D1031–D1038 (2018).

62. Kariotis, S. & Jammeh, E. BioSok/spectral_clustering_of_IPAH: v1.0.1. (2021). doi:10.5281/zenodo.5549872.

63. Pan, H. et al. Integrative Multi-Omics database (iMOMdb) of Asian Pregnant Women. Hum. Mol. Genet. (2022) doi:10.1093/hmg/ddac079.

64. Tibshirani, R. & Walther, G. Cluster Validation by Prediction Strength. J. Comput. Graph. Stat. 14, 511–528 (2005).

