## Supplementary file for "Omada: Robust clustering of transcriptomes through multiple testing"

### Omada: An unsupervised machine learning toolkit for automated sample clustering of gene expression profiles

Kariotis *et al*

#### Supplementary results

To determine the significance of the difference between simulated distributions, used during the dataset simulation, the Kolmogorov's D statistics are shown in Supplementary Table 1.

#### Supplementary results

##### Datasets

All results generated from our tools application on the available datasets can be found in Supplementary Table 2 (simulated single-class dataset), Supplementary Table 4 (simulated multi-class dataset), Supplementary Table 5 (pancan dataset), Supplementary Table 7 (I/PAH dataset) and Supplementary Table 8 (GUSTO dataset). Additionally, the confusion matrix for the simulated multi-class dataset, for which we know the actual labels, are presented in Supplementary Table 6.

#### Supplementary Tables

**Supplementary Table 1 |** Kolmogorov's D statistic for a simulated dataset containing 5 clusters (A, B, C, D, E). Each distance D has a p-value of less than 2.2e-16

| **AvsB** | **AvsC** | **AvsD** | **AvsE** | **BvsC** | **BvsD** | **BvsE** | **CvsD** | **CvsE** | **DvsE** |
|---|---|---|---|---|---|---|---|---|---|
| 0.99 | 0.99 | 1 | 1 | 0.84 | 0.97 | 0.97 | 0.65 | 0.88 | 0.54 |

**Supplementary Table 2 |** Simulated single-class dataset results, composed of 100 samples and 100 genes drawn from a single distribution

| **Tool** | **Parameters** | **Results** |
|---|---|---|
| Dataset clustering feasibility | 100 samples, 100 genes | max stability = 0.55<br>average stability = 0.45 |

| | | |
|---|---|---|
| Clustering method selection | max clusters = 6<br>comparisons = 3 | Spectral PA = **0.52**<br>Kmeans PA = 0.03<br>Hierarchical PA = 0.26 |
| Sample set selection | min(k) = 2<br>max(k) = 6<br>feature step = 20 | Optimal features = 20<br>max stability = 0.51<br>average stability = 0.47 |
| K estimation | min(k) = 2<br>max(k) = 6<br>method = spectral | Optimal k = 5 |

**max clusters: the maximum number of clusters to be tested starting from 2, range [2, max clusters]*
**min(k): minimum number of clusters to be tested*
**max(k): maximum number of clusters to be tested*
**feature step: the number of features by which the generated datasets grow. Also, the smallest dataset to be tested*

**Supplementary Table 3 |** The scores of all internal indexes used to decide on the ensemble voting of the number of clusters for the multi and single class simulated datasets. The scores for the most voted k are presented for each dataset along with the ideal score (min/max) for each index

| | **Multi Class (k=5)** | **One Class (k=6)** | **Ideal** |
|---|---|---|---|
| **Calinski-Harabasz** | 608.5824 | 5.325228 | max |
| **Dunn** | 0.672101 | 0.373477 | max |
| **Pbm** | 740.0956 | 0.709396 | max |
| **Tau** | 0.50501 | 0.192052 | max |
| **Gamma** | 0.896494 | 0.335348 | max |
| **C index** | 0.01731 | 0.324603 | min |
| **Davies–Bouldin** | 0.993194 | 2.897971 | min |
| **Mcclain rao** | 0.303852 | 0.907469 | min |
| **sd_dis** | 0.233813 | 0.479863 | min |
| **Ray–Turi** | 0.301727 | 2.20407 | min |
| **g_plus** | 0.016422 | 0.108995 | min |
| **Silhouette** | 0.479414 | 0.050476 | max |
| **s_dbw** | 0 | 0 | min |
| **Compactness** | 0 | 145.1075 | max |
| **Connectivity** | 7.512624 | 6.277666 | max |

**Supplementary Table 4 |** Simulated multi-class dataset results, where distribution is represented by around 120 samples and 3 clusters are used as a default parameter

| Tool | Parameters | Results |
|---|---|---|
| Dataset clustering feasibility | 359 samples, 300 genes | max stability = 0.78<br>average stability = 0.72 |
| Clustering method selection | max clusters = 6<br>comparisons = 3 | Spectral PA = **0.56**<br>Kmeans PA = 0.53<br>Hierarchical PA = 0.28 |
| Sample set selection | min(k) = 2<br>max(k) = 6<br>feature step = 25 | Optimal features = 300<br>max stability = 0.84<br>average stability = 0.78 |
| K estimation | min(k) = 2<br>max(k) = 6<br>method = spectral | Optimal k = 5 |

**max clusters: the maximum number of clusters to be tested starting from 2, range [2, max clusters]*
**min(k): minimum number of clusters to be tested*
**max(k): maximum number of clusters to be tested*
**feature step: the number of features by which the generated datasets grow. Also, the smallest dataset to be tested*

**Supplementary Table 5 |** Pancan multi-tissue RNA-seq dataset results using 3 cancer classes

| Tool | Parameters | Results |
|---|---|---|
| Dataset clustering feasibility | 2244 samples, 243 genes | max stability = 1<br>average stability = 0.88 |
| Clustering method selection | max clusters = 5<br>comparisons = 3 | Spectral PA = **0.63**<br>Kmeans PA = 0.44<br>Hierarchical PA = 0.06 |
| Sample set selection | min(k) = 2<br>max(k) = 5<br>feature step = 50 | Optimal features = 243<br>max stability = 0.96<br>average stability = 0.85 |
| K estimation | min(k) = 2<br>max(k) = 6<br>method = spectral | Optimal k = 3 |

**max clusters: the maximum number of clusters to be tested starting from 2, range [2, max clusters]*
**min(k): minimum number of clusters to be tested*
**max(k): maximum number of clusters to be tested*
**feature step: the number of features by which the generated datasets grow. Also, the smallest dataset to be tested*

**Supplementary Table 6 |** The confusion matrix of cluster estimations and actual classes of samples for the simulated multi-class

| | **Simulated multi-class dataset** | | | | |
|---|---|---|---|---|---|
| | **Class 1** | **Class 2** | **Class 3** | **Class 4** | **Class 5** |
| **Cluster 1** | 72 | 0 | 0 | 0 | 0 |

| | | | | | |
|---|---|---|---|---|---|
| **Cluster 2** | 0 | 0 | 0 | 72 | 0 |
| **Cluster 3** | 0 | 0 | 72 | 0 | 0 |
| **Cluster 4** | 0 | 0 | 0 | 0 | 72 |
| **Cluster 5** | 0 | 71 | 0 | 0 | 0 |

**Supplementary Table 7 |** RNA-seq iPAH/HPAH dataset results as shown in [7]

| Tool | Parameters | Results |
|---|---|---|
| Dataset clustering feasibility | 359 samples, 25955 features | max stability = 0.74<br>average stability = 0.61 |
| Clustering method selection | max clusters = 10<br>comparisons = 3 | Spectral PA = **0.86**<br>Kmeans PA = 0.17<br>Hierarchical PA = 0.57 |
| Sample set selection | min(k) = 2<br>max(k) = 10<br>feature step = 50 | Optimal features = 300<br>max stability = 0.73<br>Average stability = 0.61 |
| K estimation | min(k) = 2<br>max(k) = 10<br>method = spectral | Optimal k = 5 |

**max clusters: the maximum number of clusters to be tested starting from 2, range [2, max clusters]*
**min(k): minimum number of clusters to be tested*
**max(k): maximum number of clusters to be tested*
**feature step: the number of features by which the generated datasets grow. Also, the smallest dataset to be tested*

**Supplementary Table 8 |** Gestational diabetes dataset (GUSTO) results

| Step | Parameters | Results |
|---|---|---|
| Dataset clustering feasibility | 238 samples, 24,070 features | max stability = 0.59<br>average stability = 0.56 |
| Clustering method selection | max clusters = 5<br>comparisons = 3 | Spectral PA = **0.61**<br>Kmeans PA = 0.60<br>Hierarchical PA = 0.12 |
| Sample set selection | min(k) = 2<br>max(k) = 6<br>feature step = 50 | Optimal features = 50<br>max stability = 0.71<br>average stability = 0.65 |
| K estimation | min(k) = 2<br>max(k) = 6<br>method = spectral | Optimal k = 2 |

**max clusters: the maximum number of clusters to be tested starting from 2, range [2, max clusters]*
**min(k): minimum number of clusters to be tested*
**max(k): maximum number of clusters to be tested*
**feature step: the number of features by which the generated datasets grow. Also, the smallest dataset to be tested*

**Supplementary Table 9** | P-values of chi-square statistical and generalised linear regression analysis results for clinical variables and maternal phenotypes for the GUSTO dataset

| | Method | p-value |
|---|---|---|
| Hospital (2 locations) | chi-square | 0.072 |
| Time before/after 11am | chi-square | 0.351 |
| Sequenced location (2 locations) | chi-square | 0.468 |
| Sequenced machine (7 machines) | chi-square | 1.39e-03 |
| Sequenced Flow (22 flows) | chi-square | 2.55e-06 |
| Ethnicity (Chinese/Malay/Indian) | chi-square | 0.15 |
| Full/pre term | chi-square | 0.53 |
| Mother GDM | chi-square | 1 |
| Infant sex | chi-square | 0.119 |
| Pre-eclampsia | chi-square | 0.153 |
| ppBMI (under/normal/overweight) | chi-square | 0.235 |
| bookingBMI, WHO class(under/normal/overweight) | chi-square | 0.235 |
| Total GWG IOM (ppBMI) (Inadequate/Normal/Excessive) | chi-square | 0.577 |
| Total GWG IOM (bookBMI) (Inadequate/Normal/Excessive) | chi-square | 0.063 |
| Rate of GWG IOM (ppBMI) (Inadequate/Normal/Excessive) | chi-square | 0.311 |
| Rate of GWG IOM (bookBMI) (Inadequate/Normal/Excessive) | chi-square | 0.425 |
| Average maternal age | GLM | 0.299 |
| Average gestational weeks | GLM | 0.604 |
| Average ppBMI | GLM | 0.379 |
| Average bookingpBMI | GLM | 0.539 |
| Average total GWG (ppweight) | GLM | 0.655 |
| Average total GWG (bookingweight) | GLM | 0.258 |
| Average rate of GWG | GLM | 0.294 |
| Average EPDS | GLM | 0.602 |
| Average STAI state | GLM | 0.25 |
| Average STAI trait | GLM | 0.195 |
| Average fasting glucose | GLM | 0.696 |
| Average 2hr post glucose | GLM | 0.762 |

#### Supplementary Figures

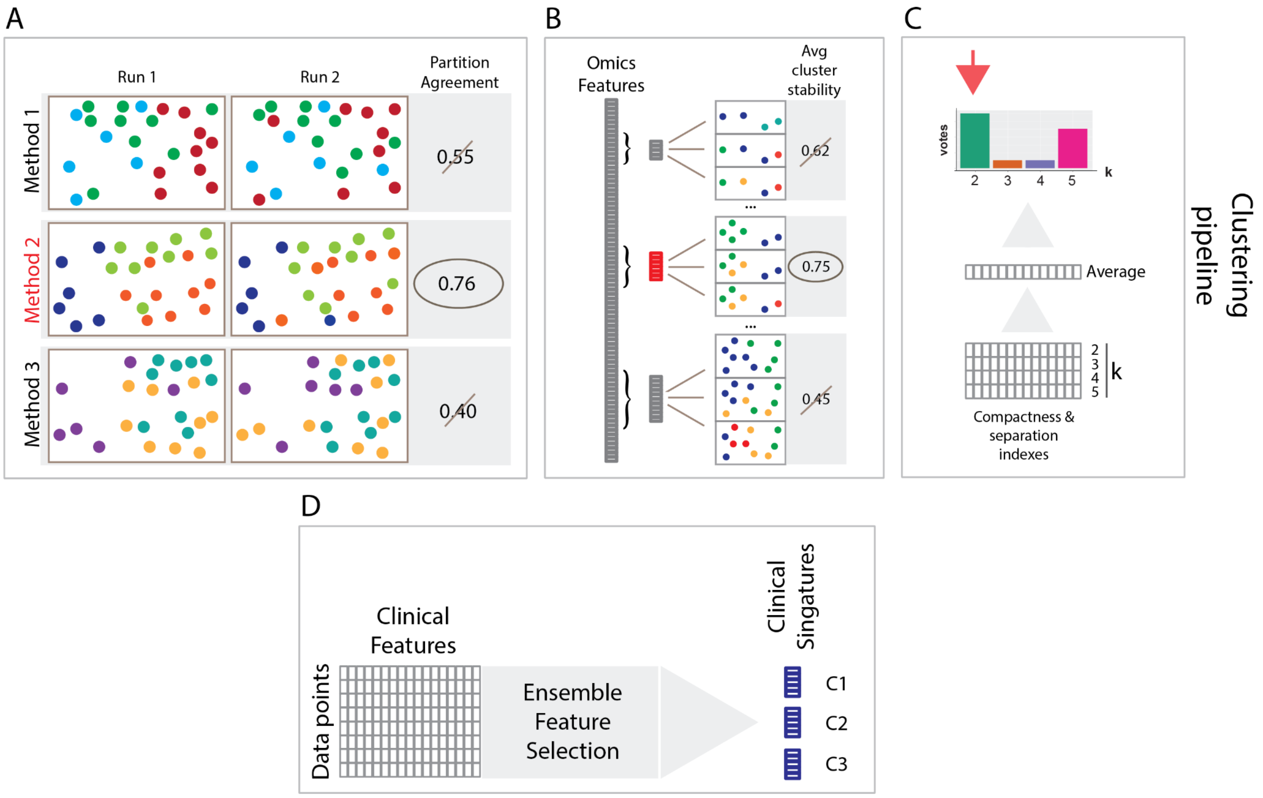


*Supplementary Figure 1: A) Clustering method selection based on the highest partition agreement between multiple differently parameterized runs B) Selection of the most cluster stable omics feature subset C) Estimating the optimal number of clusters for a dataset based on majority voting of several compactness and separation internal machine learning indexes D) Generating signature groups of clinical features/variables that drive each cluster by a multistep ensemble feature selection process utilising several machine learning classifiers*
